# Identification of the three zinc-binding sites on Tau protein

**DOI:** 10.1101/2021.04.23.441079

**Authors:** Romain La Rocca, Philipp O. Tsvetkov, Andrey V. Golovin, Diane Allegro, Pascale Barbier, Soazig Malesinski, Françoise Guerlesquin, François Devred

## Abstract

Tau protein has been extensively studied due to its key roles in microtubular cytoskeleton regulation and in the formation of aggregates found in some neurodegenerative diseases. Recently it has been shown that zinc is able to induce tau aggregation by interacting with several binding sites. However, the precise location of these sites and the molecular mechanism of zinc-induced aggregation remain unknown. Here we used Nuclear Magnetic Resonance (NMR) to identify zinc binding sites on hTau40 isoform. These experiments revealed three distinct zinc binding sites on tau, located in the N-terminal part (H14, H32, H94, and H121), the repeat region (H299, C322, H329 and H330) and the C-terminal part (H362, H374, H388 and H407). Further analysis enabled us to show that the C-terminal and the N-terminal sites are independent of each other. Using molecular simulations, we modeled the structure of each site in a complex with zinc. Given the clinical importance of zinc in tau aggregation, our findings pave the way for designing potential therapies for tauopathies.

**Highlights:**

- Zinc is known to induce tau aggregation in neurodegenerative diseases
- Zinc binding locations and mechanism are not yet clear
- Using NMR we localized 3 zinc binding site on tau
- By molecular simulations, we proposed a modeled structure of each site
- Our findings pave the way for designing potential therapies for tauopathies

## 1. Introduction

Microtubule-associated protein tau plays a central role in cytoskeleton dynamicity via its regulatory activity on microtubule dynamics, which impacts microtubule functions in various processes such as cell division and axonal stability in neurons [1,2]. In addition, tau has also been shown to be associated with a number of neurodegenerative diseases such as Alzheimer’s disease (AD), Parkinson’s disease (PD) and frontotemporal lobar degeneration, also called Tauopathies [3–6]. In several Tauopathies, tau forms neurofibrillary tangles, which consist of stacked paired helical filaments (PHF) of hyper-phosphorylated tau molecules [7].

Among the endogenous factors that have been shown to favor tau aggregation, zinc seems to play an important role in AD (see [8] for review), similarly to its role in aggregation of other proteins implicated in neurodegeneration. Indeed, zinc has been identified as a factor favoring the aggregation of amyloid-β [9,10], FUS/TLS [11], TDP-43 [12,13] and tau [14]. Zinc has been shown to increase the toxicity of tau in cells [15] mainly through an interaction with tau. More recently, this effect was confirmed in neurons where zinc induces cell death through the acceleration of tau aggregation [16,17]. Several molecular works proved that zinc can also accelerate aggregation of purified tau constructs [14,18,19], thus confirming the direct effect of zinc on tau. In addition, zinc has been recently shown to induce the formation of reversible aggregation of tau [20,21], and to promote Liquid-Liquid Phase Separation (LLPS) of tau [22,23] which could either be part of the early stages of pathological aggregation similarly to tau phosphorylation [24], or a physiological concurrent pathway.

Despite the importance of zinc binding to tau, the exact binding sites are not currently known. Two reports have already addressed this question and spotted seven residues located in the R2-R3 repeat region of tau (C291, C322, H268, H299, H329, H330, and H362) [14,16]. Among these amino acids, the cysteines located in the repeat region of tau (C291 and C322) were identified as the core of the interaction between a tau peptide (244-372) and zinc (Fig. 1). Two other studies highlighted the binding of zinc in the repeat region using the R3 repeat only [17] and a tau construct lacking the R2 repeat [18]. Two other recent studies conducted by isothermal titration calorimetry and mass spectrometry suggested presence of additional lower-affinity sites on tau [19,25]. However, the exact nature and location of these elusive sites remained unknown.

**Figure 1:**
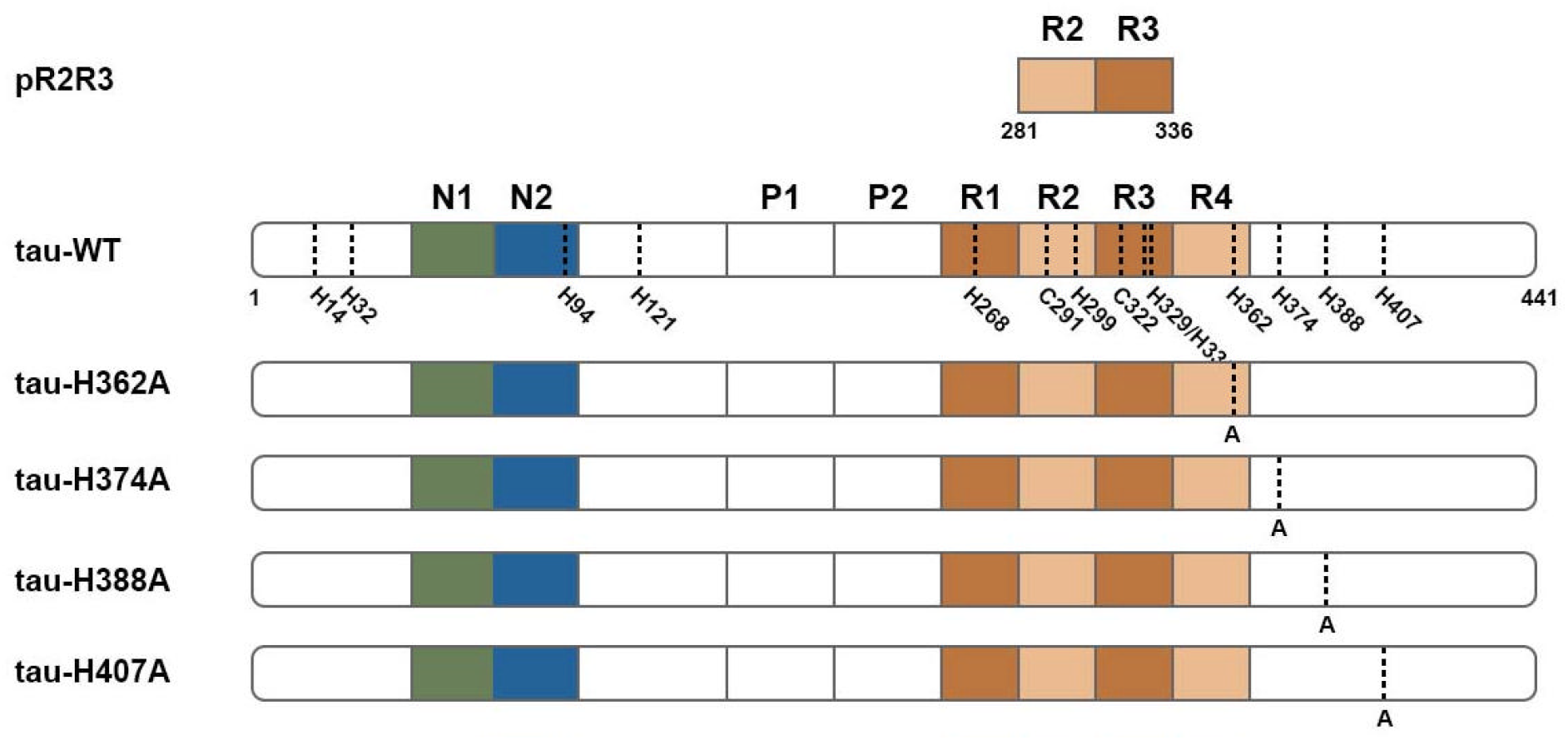
pR2R3 peptide and tau sequences of mutants. Cysteines and histidines are represented on the tau-WT sequence, and alanines mutations for each mutant are shown below histidines positions.

In this work we used a combination of Nuclear Magnetic Resonance (NMR) assays and molecular simulations to identify and propose a structure for three zinc binding sites on tau protein. We pinpoint the zinc-binding residues in the high affinity binding site in the R2R3 region and uncover the existence of two novel lower affinity binding sites in N and C terminal parts of tau.

## 2. Materials and methods

### Peptide synthesis

Synthetic pR2R3 peptide was purchased from GeneCust.

#### Protein expression and purification

The full-length tau and histidine mutants were expressed from a pET-3d vector introduced into Escherichia coli BL21(DE3). After 3 h of induction with 0.5 mM isopropyl β-D-1-thiogalactopyranoside (IPTG), cells were centrifuged and the pellets were resuspended into a lysis buffer as previously described [26]. Lysis was pursued using three runs of French press at 4 tones, and non-thermostable proteins were precipitated at 95°C for 11 minutes. Then, the lysate was centrifuged at 30 000 g for 30 min at 4°C, and the supernatant was injected onto a HiTrap SP Sepharose HP cation exchange column pre-equilibrated with a 50 mM MES pH 6.5 buffer. Tau was eluted with a 50 mM MES NaCl 0.5 M pH 6.5 buffer at 1 mL/min, dialysed three times against 2 L of water at 4°C for 2 h, and dry-lyophilized. For ^15^N labeled-tau, cells were grown on LB medium, centrifuged and the pellet was solubilized in M9 medium containing ^15^N ammonium chloride. Prior to use, each tau sample was airfuged at 25 psi and dosed at 280 nm with an extinction coefficient of 7700 M^-1^.cm^-1^ by spectrophotometry as previously described [26].

#### NMR spectroscopy

NMR spectra were recorded on a Bruker spectrometer 600 MHz equipped with a TCI cryoprobe (Triple Resonance Probe). NMR experiments (^15^N-HSQC) were performed at 280°K (7°C) on 50 μM for ^15^N-labeled tau samples and 1 mM for pR2R3 peptide. For the pH variations experiments, tau was resuspended into a sodium phosphate buffer at pH 6.1 and 6.7 respectively. For zinc titrations, the buffer was 20 mM MES, 100 mM NaCl, pH 6.5. Using previous chemical shift assignments, we identified the cross-peaks corresponding to the N-terminal and C-terminal extremities in the full-length tau (BMRB entry 17920 [27]) and repeat region in the pR2R3 spectrum (BMRB entry 19253 [28]). H362, H374, H388 and H407 were assigned on the tau-WT spectrum by using mutants of tau. Data was processed and analyzed with TopSpin software from Bruker.

#### Molecular simulations

The Rosetta *de novo* structure prediction method with Monte-Carlo approach was utilized for structure prediction for tau fragments with zinc coordination restraints. Fragment library was built with the Robetta web server for the whole tau protein isoform hTau40 (UniProt identifier : P10636-8). This fragment library was adopted to three selected regions of tau. Simulations were started from an extended chain and then subjected to *ab initio* folding protocol with zinc coordination restraints with scoring function ref2015. Zinc incorporation was done as it was described in the original report with distance and angle restraints to the supposed coordination sphere. All backbone and sidechain torsions and zinc rigid-body degrees of freedom were optimized simultaneously in the all-atom refinement stage according to the original report [29]. For each of three Tau regions, 45,000 models were generated and the first 100 models were selected for further analysis. The top model was chosen based on the pair score.

## 3. Results

### 3.1. Localization of zinc-binding sites

Many studies have concluded that the amino acids responsible for zinc binding to tau are located in the R2R3 repeat region of tau [14,15,17,18]. More recently it was suggested that additional sites would exist most likely located outside of this repeat region [19,30]. In order to identify the zinc binding sites on tau protein, we performed ^15^N-Heteronuclear Single Quantum Coherence (HSQC) NMR on tau first in the absence, and then in the presence of zinc. Since the amino acid assignment is not complete for full length tau [27], based on the existing attributions, we conducted two separate sets of NMR experiments, first on the peptide pR2R3, to identify the amino acids implicated in zinc binding in the repeat region, and second on full-length tau, to identify the amino acids implicated in zinc binding outside of the repeat region (see Fig. S1).

#### 3.1.1. Identification of zinc binding residues in R2R3 region of tau

To first explore the R2R3 repeat domain of tau, we recorded the ^15^N-HSQC spectrum at ^15^N-natural abundance on pR2R3 peptide (pR2R3, Figure 1) in the absence of zinc (Figure 2A, orange cross peaks). Using the assignment already published for this region (Biological Magnetic Resonance Bank, BMRB, entry 19253 [28]), we identified cross-peaks corresponding to all Cys and His residues in this region (C291, H299, C322, H329 and H330). It should be noted that the cross-peak corresponding to C291 was split, suggesting the existence of two possible conformations for this residue.

**Figure 2:**
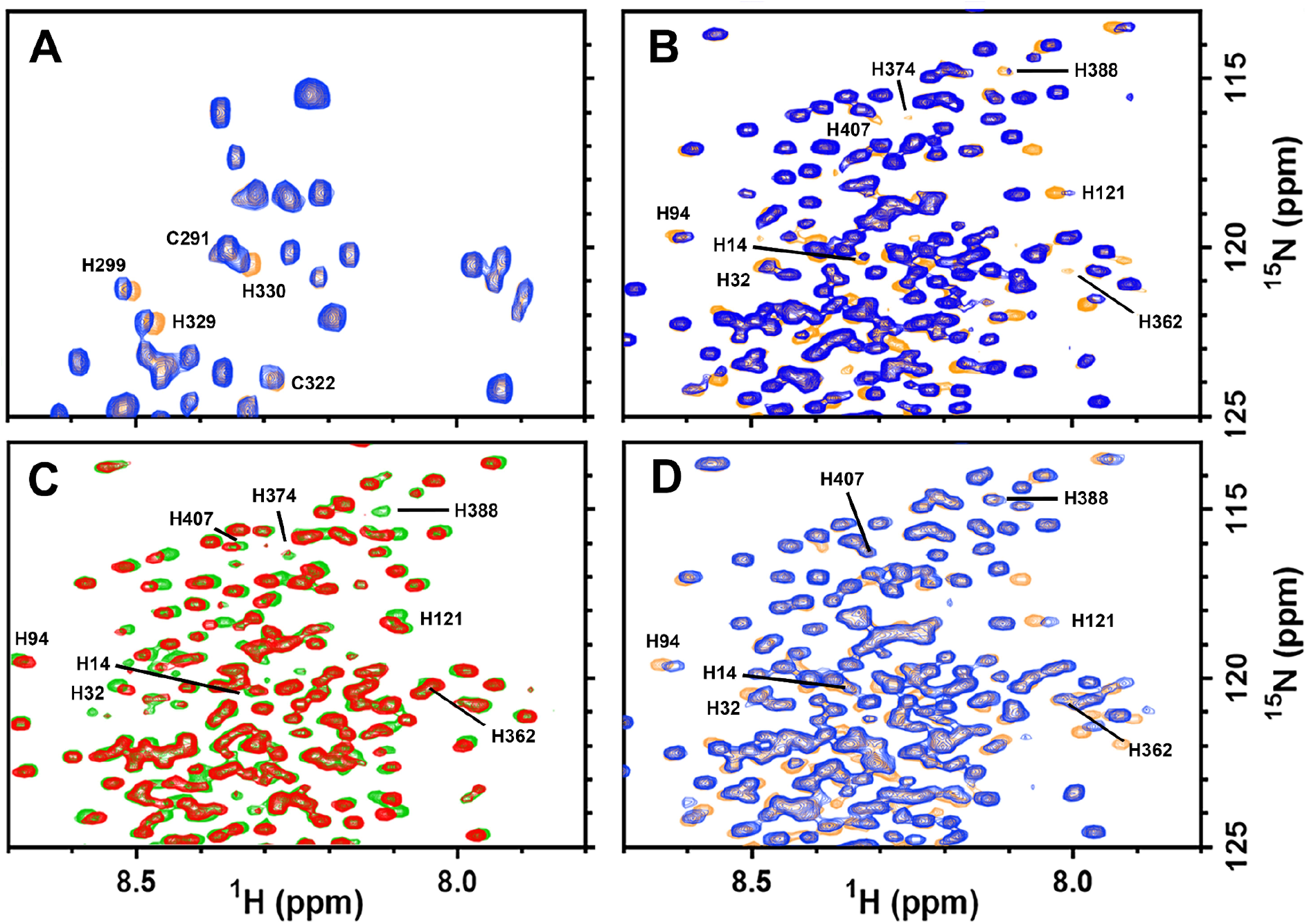
^15^N HSQC spectra of tau constructs. A: ^15^N HSQC spectrum of 1 mM unlabeled pR2R3 at natural abundance in the absence (orange peaks) or in the presence of 3 mM zinc (blue peaks). B: ^15^N HSQC spectrum of 50 μM ^15^N-labeled tau-WT in the absence (orange peaks) or in the presence of 300 μM zinc (blue peaks). C: ^15^N HSQC spectrum of 50 μM ^15^N-labeled tau-WT at pH 6.1 (green peaks) and 6.7 (red peaks). D: ^15^N HSQC spectrum of tau-H374A in the absence (orange peaks) or in the presence of zinc (blue peaks). All the experiments were done at 7°C to avoid aggregation in 20 mM MES 100 mM NaCl pH 6.5 unless otherwise stated.

In the presence of zinc (Fig. 2A, blue peaks), only the cross-peaks corresponding to H299, C322, H329 and H330 were shifted, indicating that these four residues of the R2R3 region participate in the chelation of zinc. Interestingly, the addition of zinc only slightly influenced the splitting of C291 cross peak thus possibly favoring one of the two conformations, but not contributing to zinc binding.

#### 3.1.2. Identification of zinc binding residues in N- and C-terminal regions of tau

To explore the N- and C-terminal regions of tau, first we recorded the ^15^N-HSQC spectrum of full-length tau (Fig. 1, tau-WT, wild type) in the absence of zinc (Fig. 2B, orange peaks). Using the previously published amino-acid assignment map (BMRB, entry 17920 [27]), we identified the cross-peaks corresponding to H14, H32, H94 and H121 in the N-terminal region. Since none of the four C-terminal histidines were assigned in this map, we generated single amino-acids mutants for each of the four histidines of this region (Fig. 1, tau-H362A, tau-H374A, tau-H388A, and tau-H407A) and recorded ^15^N-HSQC spectra of these mutants (see Fig. S2, orange peaks). This enabled us to identify the cross-peaks corresponding to H362, H374, H388 and H407 on the full-length tau spectrum (Fig. 2B, orange peaks).

In order to confirm that all the cross peaks we identified as histidines in the N-terminal region are indeed histidines, ^15^N-HSQC spectrum of full-length tau was recorded at two different pH (pH 6.1 and pH 6.7) to bring the pH closer to the pKa of histidine. Indeed the cross-peaks corresponding to putative histidine shifted between the two conditions (Fig. 2C, green and red peaks).

Having now identified all the histidines in the C-terminal and N-Terminal regions, we performed NMR on full-length tau in the presence of zinc (Fig. 2B, blue peaks). The addition of zinc induced a shift of cross-peaks of H14, H32, H94, H121, H362, H374, H388 and H407, which indicated that these residues participate in the chelation of zinc. As expected, cross-peaks corresponding to amino-acids in the proximity of these histidines were also shifted (see Fig. S3). Considering the localization of the histidines we identified, our results let us hypothesize the existence of two additional binding sites : one binding site in C-terminal (H362, H374, H388 and H407) and one binding site in N-terminal (H14, H32, H94, H121).

#### 3.1.3. Zinc binding sites in N- and C-terminal regions are independent

In order to demonstrate that the eight identified histidines form two separate independent sites at the N-terminal and in C-terminal of tau, we performed NMR on tau with one of its C-terminal His mutated into Ala (Fig. 1, tau-H374A) in the presence of zinc (Fig. 2D). Upon mutating this amino acid, the cross peaks corresponding to the other three histidine residues located at the C-terminus did not shift anymore after adding zinc indicating a loss of zinc binding. Furthermore, this mutation in the C-terminal did not have any impact on zinc-dependent shifting of the cross-peaks corresponding to the histidines in the N-terminal region (H14, H32, H94 and H121). Similar results were obtained with mutation of the three other C-terminal His (Fig. S2). This indicated the existence of two independent zinc binding sites, one in the C-terminal and one in the N-terminal region.

In summary, NMR experiments revealed the presence of three zinc-binding sites in tau, involving histidine and cysteine residues, with one site located in the R2R3 region, and two sites in the N-terminal and in the C-terminal region, independent of each other.

### 3.2. Modelisation of zinc binding sites of tau

NMR data allowed us to modelize the three sites identified in our study with a molecular simulation using three small tau peptides encompassing each zinc binding site. N-terminal site was modelized using Tau peptide (13-122) (Fig. 3A), R2R3 site was modelized using Tau peptide (298-331) (Fig. 3B), and the C-terminal site was modelized using Tau peptide (361-408) (Fig. 3C). In all three cases molecular modelisation revealed that the presence of zinc induced a more condensed conformation of tau by bringing together amino acids that would otherwise be distant. This compaction may result in a more globular shape of tau, which is in agreement with previous published DLS data [31].

**Figure 3:**
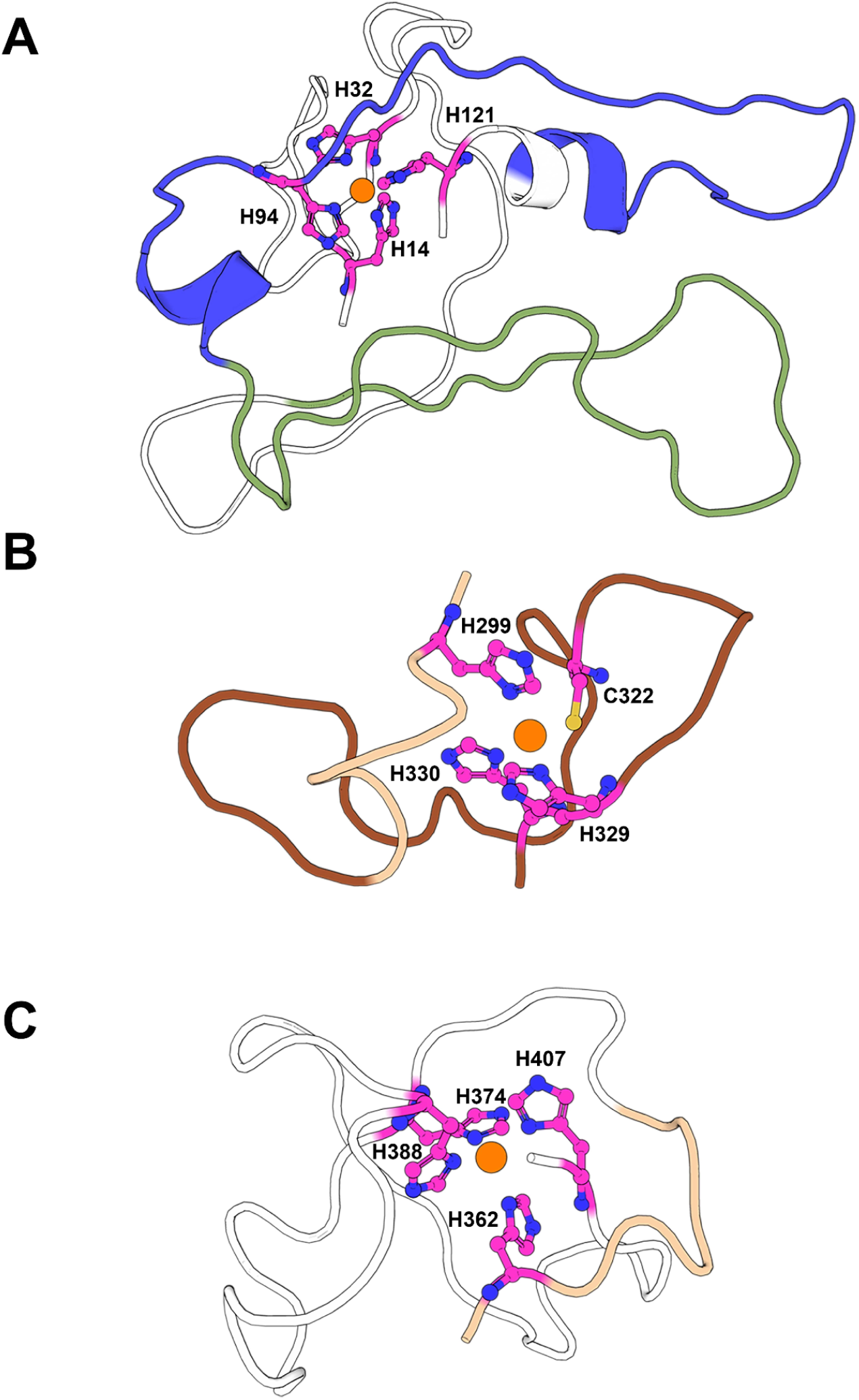
3D models of the zinc-binding sites on tau. A: N-terminal (13-122), B: R2R3 (298-331) and C: C-terminal (361-408) regions of tau in the presence of zinc (orange sphere).

## 4. Discussion

Despite the clinical importance of tau protein and the recent discovery of zinc ions influencing the aggregation and self-assembly of tau, the number of tau-zinc interaction sites was still under debate [19,32]. In this study we identify 12 zinc-binding amino acids corresponding to three zinc-binding sites (one in the R2R3 region, one in N-terminal and one in C-terminal part). Our findings are consistent with the fact that zinc is mainly tetra-coordinated in proteins by Cys and His amino-acids [33,34].

Among the six amino acids that have been previously suggested to participate in zinc-binding to the R2R3 region of tau [14,16–18], we finally pinpoint the 4 amino acids (H299, C322, H329 and H330) constituting the main zinc-binding site in tau. This site has been previously described as the high-affinity site (10^6^ M^-1^), as mutations of cysteins and histidines of these regions disrupted the high-affinity interaction between tau repeat regions and zinc [14]. The same study showed that zinc is able to greatly increase the formation of ThT-positive tau peptide fibrils. Moreover, it has been shown that zinc decreases the survival of tauopathy model cells [15] and neuronal cells [16] by interacting directly with at least one cysteine of tau. Our data allowed us to identify the four amino-acids responsible for this toxicity, thus providing the first molecular model of the zinc high-affinity binding site on tau.

In addition, we now describe two novel zinc binding sites in the C-terminal and N-terminal regions of tau. These sites may correspond to the low-affinity ones, previously observed with ITC experiments [20]. First, the C-terminal low-affinity binding site is composed of H362, H374, H388 and H407, and is the closest site from the R2R3 region. One explanation of its role may be linked to a previous work, which describes an interaction between the R3 and the R4 regions of tau preventing the aggregation of a R1R3R4 tau peptide [18]. Using NMR and aggregation assays, the authors have found that zinc can disrupt this interaction by competing with specific tau amino-acids, thus enhancing the aggregation of the peptide. Given that the C-terminal site starts at the end of the R4 region of tau, zinc could potentially trigger tau aggregation by preventing such interaction between R3 and R4 repeats. Second, the N-terminal low-affinity binding site is composed of H14, H32, H94 and H121, and is located far from the two other sites. However, similarly to the C-terminal site, it could play a role in the triggering of the aggregation process, since only high zinc concentration is able to trigger the aggregation of tau [20], thereby highlighting the role of the low-affinity sites in this process. Thus, the zinc concentration could be one major factor of the aggregation of tau, especially considering the importance of zinc homeostasis in physiopathology [8,35].

In AD for example, intraneuronal levels of zinc in cortical cells of patients have been shown to be increased in comparison to healthy cells [36]. Similarly, high concentrations of zinc were found inside the somata and dendrites of AD neurons [37]. This accumulation may be linked to the functionality of zinc-bound proteins, which can be impaired due to alteration of neuronal metabolism in pathology [35,38,39]. Importance of zinc binding to tau has also been evoked in a physiological context, when the zinc concentrations are lower [40]. Thus, zinc binding to tau may have physiological implications with low-intraneuronal concentrations of zinc, while possibly affecting the pathological aggregation found in tauopathies at high zinc concentration. The 3D models of the R2R3 site, N-terminal site and C-terminal site that we described here provide a molecular basis for future tau functionality and tau aggregation investigations, in order to decipher this mechanism that has multiple implications in neurophysiopathology.

## 5. Conclusion

In this study, we identify three zinc binding sites of tau protein and provide a molecular model of zinc ions binding on tau. Overall, the molecular basis of the tau-zinc interaction provided in our study will help understanding the zinc-related tau regulation or aggregation processes in both physiological and pathological conditions, thus paving the way for designing potential therapies for tauopathies.

## Acknowledgements

NMR experiments were conducted in the NMR platform of IMM, Marseille, France. Molecular modelization at shared research facilities of HPC computing resources at Lomonosov Moscow State University.

