## Supplementary Data for "Identification of the three zinc-binding sites on Tau protein"

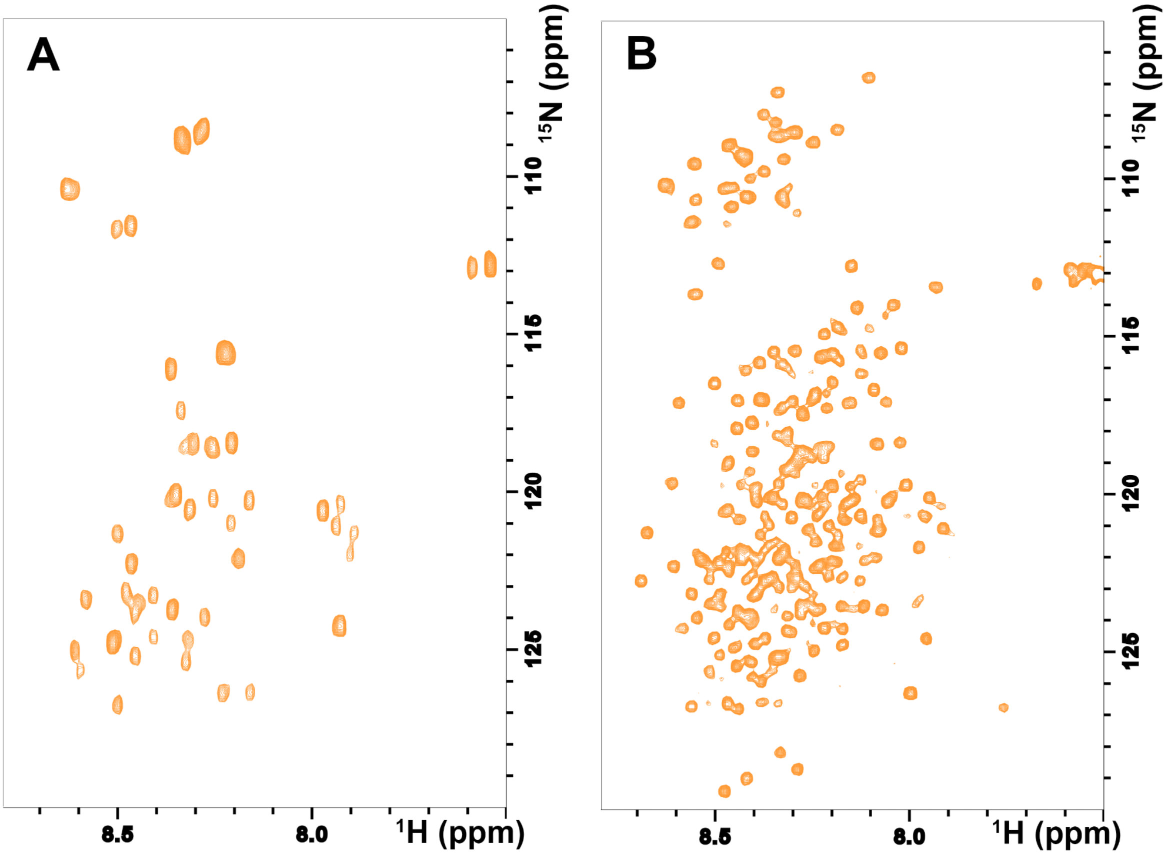


**Figure S1 : $^{15}$N HSQC NMR spectra of pR2R3 peptide (A) and tau-WT (B).** Both of the spectra were recorded at 7°C in 20 mM MES 100 mM NaCl at pH 6.5.

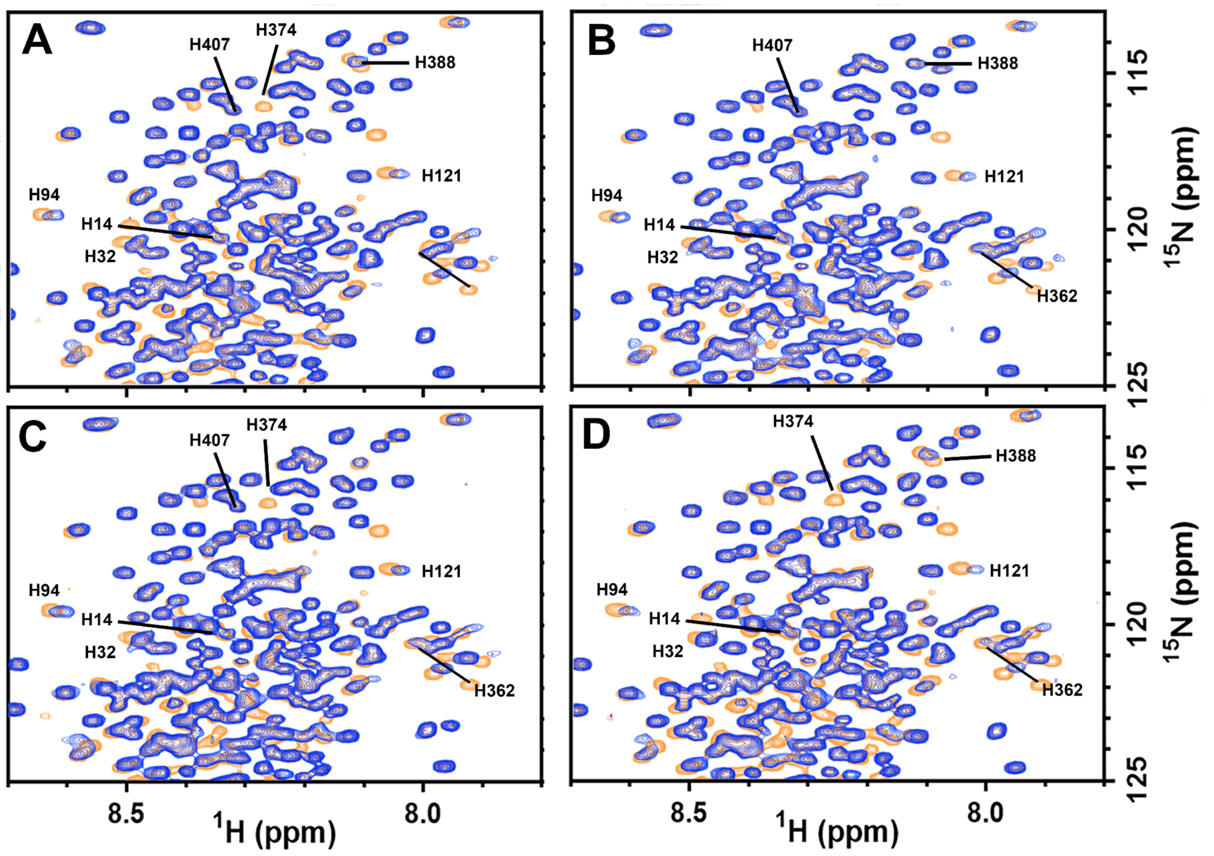


**Figure S2: $^{15}$N HSQC spectra of tau histidines mutants.** $^{15}$N HSQC spectra of tau-H362A (A), tau-H374A (B) (replicate of Figure 2D), tau-H388A (C), and tau-H407A (D) in the absence (orange) or in the presence of zinc (blue). 50 µM of $^{15}$N-labeled tau-H362A, tau-H374A, tau-H388A and tau-H407A were titrated with 300 µM of zinc at 7°C. N-terminal and C-terminal histidines are shown in cyan. All of the N-terminal histidines (H14, H32, H94 and H121) were not affected by any mutation in all spectra, as they were still shifted upon the addition of zinc. C-terminal histidines were affected differently for each mutant, underlying two independent sites at the N-terminal and C-terminal regions of tau.

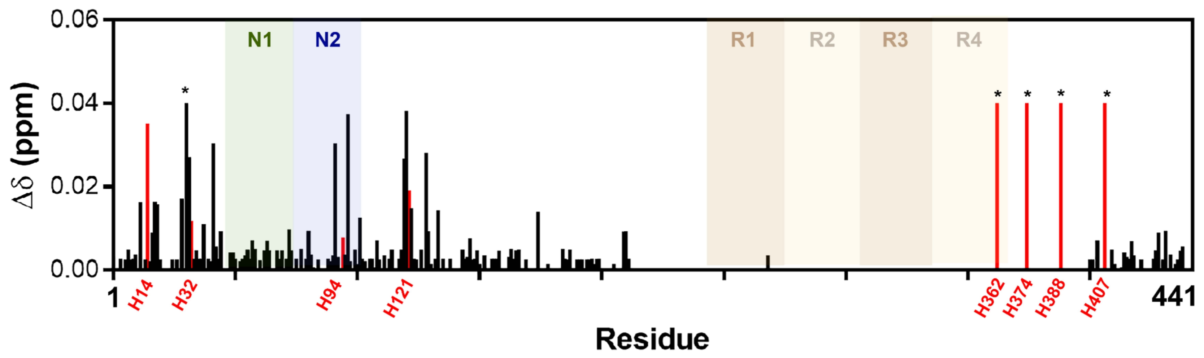


**Figure S3: Shifts of each tau-WT assigned peak in the spectrum upon the addition of zinc.** The residues annotated with * did not shift but disappeared from the spectrum upon the addition of zinc, thus indicating a high perturbation.
